# A user’s guide to the online resources for data exploration, visualization, and discovery for the Pan-Cancer Analysis of Whole Genomes project (PCAWG)

**DOI:** 10.1101/163907

**Authors:** Mary Goldman, Junjun Zhang, Nuno A. Fonseca, Isidro Cortés-Ciriano, Qian Xiang, Brian Craft, Elena Piñeiro-Yáñez, Brian D O’Connor, Wojciech Bazant, Elisabet Barrera, Alfonso Muñoz-Pomer, Robert Petryszak, Anja Füllgrabe, Fatima Al-Shahrour, Maria Keays, David Haussler, John N. Weinstein, Wolfgang Huber, Alfonso Valencia, Peter J. Park, Irene Papatheodorou, Jingchun Zhu, Vincent Ferretti, Miguel Vazquez, on behalf of the PCAWG Portals and Visualization Working Group, the ICGC/TCGA Pan-Cancer Analysis of Whole Genomes Network

**Affiliations:** UC Santa Cruz Genomics Institute, Santa Cruz, 95064, USA; Ontario Institute for Cancer Research, Toronto, ON M5G 0A3, Canada; European Molecular Biology Laboratory, European Bioinformatics Institute, EMBL-EBI, Hinxton, CB10 1SD, UK; Department of Biomedical Informatics, Harvard Medical School, Boston, MA, USA; Centre for Molecular Science Informatics, Department of Chemistry, University of Cambridge, Lensfield Road, Cambridge CB2 1EW, United Kingdom; Bioinformatics Unit, Spanish National Cancer Research Centre (CNIO), Madrid, 28029, Spain; UT MD Anderson Cancer Center, Houston, 77030, USA; European Molecular Biology Laboratory, Heidelberg, 69117, Germany; Barcelona Supercomputing Center (BSC), Barcelona, 08034, Spain; ICREA, Barcelona, 08010, Spain; CHU Sainte-Justine Research Center, Montreal, Quebec, H3T 1C5, Canada; Norwegian University of Science and Technology, Trondheim, Norway

## Abstract

The Pan-Cancer Analysis of Whole Genomes (PCAWG) project has generated, to our knowledge, the largest whole-genome cancer sequencing resource to date. Here we provide a user’s guide to the five publicly available online data exploration and visualization tools introduced in the PCAWG marker paper: The ICGC Data Portal, UCSC Xena, Expression Atlas, PCAWG-Scout, and Chromothripsis Explorer. We detail use cases and analyses for each tool, show how they incorporate outside resources from the larger genomics ecosystem, as well as demonstrate how the tools can be used together to more deeply understand tumor biology. Together, these tools enable researchers to dynamically query complex genomics data and integrate external information, enabling and enhancing PCAWG data interpretation. More information on these tools and their capabilities is available from The PCAWG Data Portals and Visualizations Page (http://docs.icgc.org/pcawg).

## Main text

The Pan-Cancer Analysis of Whole Genomes (PCAWG) project provides, to our knowledge, the largest uniformly analyzed, publicly available whole-genome dataset in cancer genomics. The PCAWG project has reprocessed whole-genome sequencing data for 2,658 donors from 48 cancer projects across 47 different primary tumor types (Campbell 2020). Here we provide a user guide to five tools introduced in the PCAWG marker paper: The ICGC Data Portal, UCSC Xena, Chromothripsis Explorer, Expression Atlas, and PCAWG-Scout, each of which was created or expanded to explore this resource (Campbell 2020). These tools aim to make PCAWG data easy to visualize and analyze by pre-loading the PCAWG data so that users do not need to locate or manage data, and by making the tool accessible through a web interface so that no extra software is needed. Each of these five tools has also focused on integrating other genomics datasets and tools that help provide insights into the genomic patterns within the PCAWG data as newly generated cancer datasets can only fully realize their potential when integrated with other genomics resources. Some of the datasets and tools integrated include the UCSC Genome Browser (Kent 2002), Ensembl (Zerbino 2018), drug target compendia (Piñeiro-Yáñez 2018), COSMIC (Shepard 2011), and even large sequencing efforts such as GTEx (Carithers 2015). Intuitive access to these additional tools and datasets is provided either by showing their data side-by-side or providing context-dependent URL links.

The five resources covered in this paper provide a different perspective and focus to the PCAWG data (Table 1). The ICGC Data Portal serves as the main entry point for accessing all PCAWG data and can also be used to explore PCAWG consensus simple somatic mutations, including point mutations and small indels, each by their frequencies, patterns of co-occurrence, mutual exclusivity, and functional associations. UCSC Xena integrates diverse types of genomic and phenotypic/clinical information at the sample-level across large number of samples, enabling rapid examination of patterns within and across data types. The Chromothripsis Explorer visualizes genome-wide mutational patterns, with a focus on complex genomic events, e.g., chromothripsis and kataegis. This is done through interactive Circos plots for each tumor with different tracks corresponding to allelespecific copy number variants, somatic structural variations, simple somatic mutations, indels, and clinical information. The Expression Atlas focuses on RNA-seq data, supporting queries in either a baseline context (e.g., find genes that are expressed in prostate adenocarcinoma samples) or in a differential context (e.g., find genes that are under-or over-expressed in prostate adenocarcinomas compared to “adjacent-normal” prostate samples). PCAWG-Scout allows users to run their own analysis on-demand, including predicting cancer driver genes, differential gene expression, calling recurrent structural variations, survival analysis, pathway enrichment, visualization of mutations on a protein structure, analysis of mutational signatures, and predictions of recommended therapies based on their in-house resource, PanDrugs (Supplementary Figure 1). Each of these five tools offers different visualizations and analyses of the PCAWG data resource, each with their strengths, and each enabling different insights into the data. When used together they provide users a deeper understanding of the tumor’s biology (Figure 1). Researchers interested in these tools can learn about and access them at the PCAWG Landing Page (http://docs.icgc.org/pcawg).

**Figure 1.**
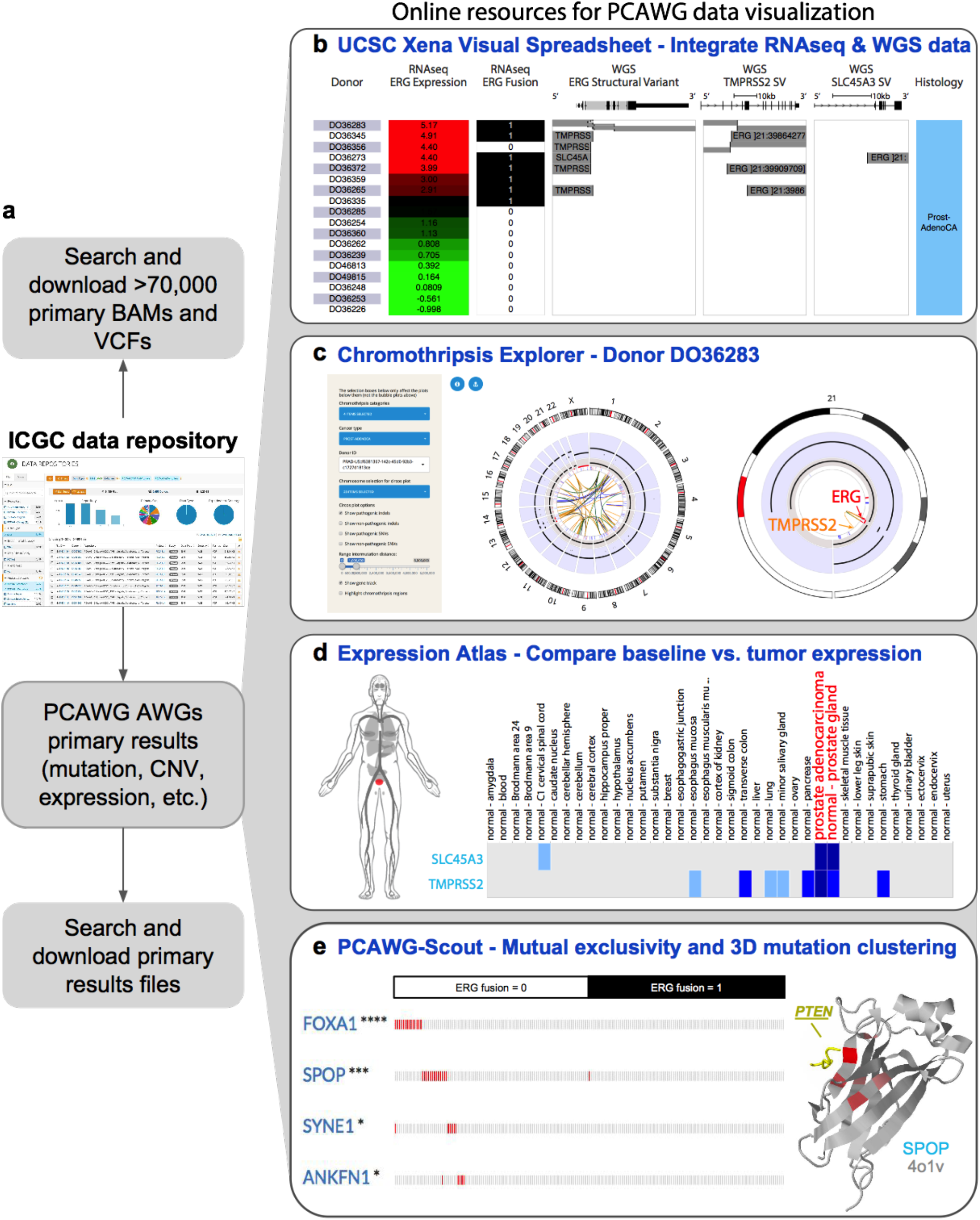
Combining strengths from each tool. These resources provide complementary views and analyses, together enabling users to gain insights into tumor biology. This is illustrated by the example of mutually exclusive, recurrent events (such as *ERG* fusions and mutations in *SPOP*) detected in prostate adenocarcinoma. **(a)** All PCAWG BAMs, VCFs, and Analysis Working Group (AWG) primary results files are available through the ICGC data portal’s Data Repository tool (https://tinyurl.com/yburnde5). To obtain these files, the user downloads a manifest of selected files and then downloads the actual data files (with authorization if needed) using the ICGC download tool. UCSC Xena, Chromothripsis Explorer, Expression Atlas and PCAWG-Scout have each pre-processed the same primary analysis working group results files. **(b)** The *UCSC Xena Visual Spreadsheet* shows that the *ERG* fusion is present in 8 out of 18 PCAWG prostate adenocarcinoma samples (https://tinyurl.com/y78adbl5), as detected by the PCAWG RNA-seq and wholegenome sequencing data. Each row corresponds to a sample. Columns, starting at the left, correspond to histology, *ERG* gene expression, and *ERG* fusion based on RNA-seq data. The next three columns show structural variant calls using whole genome DNA-seq data for *ERG, TMPRSS2*, and *SLC45A3*. **(c)***Chromothripsis Explorer* provides an in-depth genome-wide view of the copy number alterations and structural variations in the 8 tumors with *ERG* fusion listed in b. Details about the total and minor copy number, as well as the SVs, can be obtained by hovering over these elements in the interactive version of the Circos plot. Circos plot visualizations for the other 7 donors are given in Supplementary Figure 6. **(d)** The *Expression Atlas* shows a heatmap of genes (rows) and tissue or disease type (columns). Here we show the expression of *ERG, SLC45A3*, and *TMPRSS2* in healthy human tissue (top heatmap), as derived from our re-analysis of the GTEx data set. The bottom heatmap shows expression in PCAWG data (https://tinyurl.com/y9fefymf). The human figure, also known as an anatomogram, shows the highlighted prostate tissue. **(e)** *PCAWG-Scout* complements the above analysis by identifying recurrent mutational events in tumors without ERG fusion (fusion = 0). On the left, PCAWG-Scout shows a mutation exclusivity analysis (using Fisher’s exact test, n=199), which identifies *FOXA1* (****, < 0.0001), *SPOP* (***, < 0.001), *SYNE1* (*, < 0.05), and *ANKFN1* (*, < 0.05) as significantly associated with non-fusion tumors (https://tinyurl.com/ybqpou52). In the 3D protein structure of SPOP shown on the right, mutations are seen to cluster tightly around the region that overlaps with the interaction surface of *PTEN*. The portion of *PTEN* that interacts with SPOP is shown in yellow, along with the SPOP structure. Red indicates recurrent mutations in *SPOP* with a brighter red indicating higher rate of recurrence.

**Table 1.** Search, visualization, analysis/integration, and download functionalities provided by each of the PCAWG data resources.

| Functionality |  | ICGC Data Portal | UCSC Xena | Chromothripsis Explorer | Expression Atlas | PCAWG-Scout |
| --- | --- | --- | --- | --- | --- | --- |
| Search | Search by demographic data, specimen phenotype, molecular subtype | Y | Y |  |  | Y |
|  | Search by genes and/or variants | Y | Y |  | Y | Y |
|  | Search by genomic coordinates | Y | Y |  |  |  |
| Visualize | Visualize multiple types of data together |  | Y | Y | Y | Y |
|  | Visualize coding variants | Y | Y | Y |  | Y |
|  | Visualize non-coding variants | Y | Y | Y |  |  |
|  | Visualize structural variants |  | Y | Y |  | Y |
|  | Visualize mutational signatures and predicted drivers |  |  |  |  | Y |
|  | Visualize genome-wide profiles, including LOH, in Circos plots |  |  | Y |  |  |
|  | Visualize tissue expression on a human figure |  |  |  | Y |  |
|  | Visualize gene co-expression |  | Y |  | Y |  |
|  | Visualize pathways, therapeutic associations | Y |  |  |  | Y |
|  | Visualize summary of BAMs/VCFs | Y |  |  |  |  |
| Analysis | Kaplan-Meier analysis with statistics | Y | Y |  |  | Y |
|  | Geneset/pathway enrichment analysis | Y |  |  | Y | Y |
|  | View non-identifiable analysis results of protected data | Y |  | Y | Y | Y |
|  | Discover differentially- or co-expressed genes, mutually exclusive genomic events |  |  |  |  | Y |
|  | Annotations from other resources | Y | Y |  | Y | Y |
| Download | Programmatic data download | Y | Y |  | Y | Y |
|  | Download BAMs, VCFs, primary files | Y |  |  |  |  |
|  | Download secondary processed data | Y | Y |  | Y | Y |

### ICGC Data Portal and a Use Case

(https://dcc.icgc.org). As a main entry point, the ICGC Data Portal (Zhang 2019) provides an intuitive graphic interface for users to browse, search and explore PCAWG data sets (Figure 1a). Uniformly aligned sequencing BAM files and variant calling VCF files, although physically residing in multiple repositories globally, can be centrally searched via a faceted search interface (https://icgc.org/ZEA). Users can easily find specific data sets of interest with a few mouse clicks. A universal data access tool called “icgc-get” can be used to download data from different repositories. Other downstream analysis results generated by PCAWG working groups are available at https://dcc.icgc.org/releases/PCAWG. Close to 23 million open access PCAWG consensus simple somatic mutations have been annotated with consequences in protein changes, affected pathways, targeting cancer drugs and gene ontology terms alongside with clinical information integration. The portal’s Advanced Search (https://icgc.org/ZzP) tool allows users to perform complex queries, such as, retrieving the most frequently mutated genes that are targets of drugs from stage 2 liver cancer patients (https://icgc.org/ZHe). Analytic tools, including access to a Jupyter Notebook sandbox for advanced users, support exploration of potential associations between molecular abnormalities and phenotypic observations such as patient survival (https://dcc.icgc.org/analysis). The ICGC Data Portal publicly displays non-identifiable analysis results of protected data.

The ICGC Data Portal is best for users who are seeking to download PCAWG data for their own analysis. It also has the richest resources and functionality for those users interested in SNVs, including patterns of co-occurrence, mutual exclusivity, and functional associations. An example use case using the ICGC Data Portal is found in Figure 1a, demonstrating how bioinformaticians and other tool creators can download results from the portal and then run their own analyses or offer up their own visualizations of this data.

### UCSC Xena and a Use Case

(https://pcawg.xenahubs.net). UCSC Xena’s adaptable visualizations, fast performance, and flexible data format bring the full power of the PCAWG resource to all researchers (Goldman 2018). It displays data mapped to coding and non-coding regions of the genome, including introns, promoters, enhancers, and intergenic regions. Xena can display tens of thousands of data points on thousands of samples, all within seconds. Xena visualizes a comprehensive list of PCAWG primary and derived results (Campbell 2020). The Xena Browser excels at integrating the diverse datasets generated by the PCAWG Consortium using the Xena Visual Spreadsheet, which enables users to view multiple types of data side-by-side (Figure 1b). In addition to the Visual Spreadsheet, Xena offers survival analyses, the ability to compare and contrast dynamically built subgroups, statistical tests such as ANOVA and others, and URLs to live visualizations for sharing with collaborators or others. Xena’s hub-browser architecture enables users to view the protected consensus simple somatic mutations, including non-coding mutations, by loading the dataset into a user’s local private Xena hub (Figure 2, Supplementary Figure 3). The Xena Browser seamlessly integrates data from multiple hubs, allowing users who have access to the protected mutation data to visualize it in conjunction with other PCAWG data publicly available on the PCAWG Xena Hub (https://pcawg.xenahubs.net).

**Figure 2.**
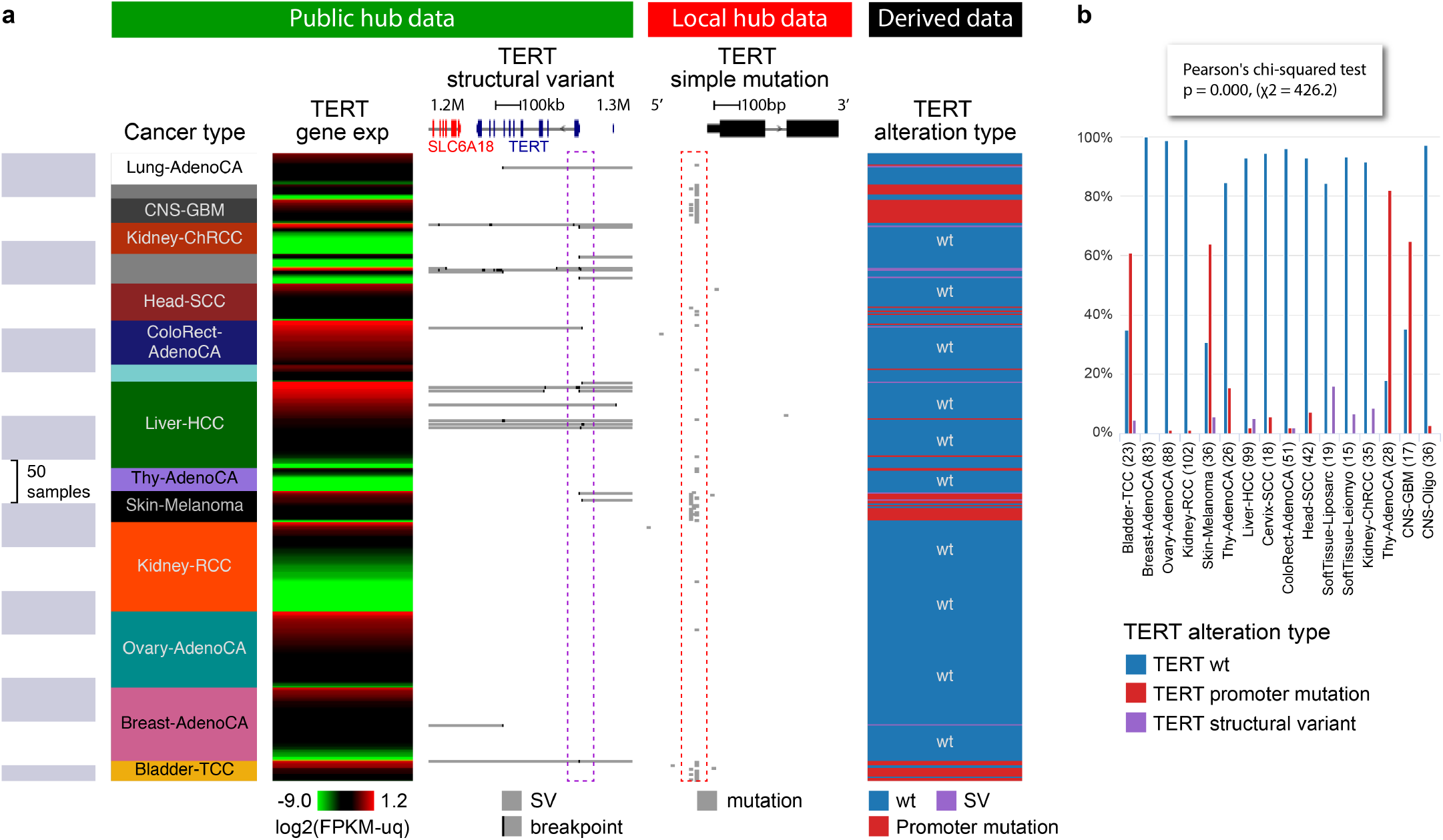
UCSC Xena views of *TERT* across cancer types. **(a)** Visual Spreadsheet view of *TERT* multi-omics data across PCAWG cancer types. Data from the PCAWG public hub are under the green section while protected data from the user’s local Xena hub is under the red section. Data is integrated in the browser, keeping private data protected. We can see that many cancer types have *TERT* alterations either as simple somatic mutations in the promoter region, as seen in the pileup highlighted in the red box, or as structural variants, as seen in the breakpoint pileup upstream of *TERT* highlighted in the purple box. Only cancer types that have a TERT alteration are displayed (n=718 samples). The last column was dynamically generated in the browser and shows which samples have promoter mutation, which have structural variants, and which have none. No sample is observed to have both promoter mutations and structural variants, as these two types of alterations are mutually exclusive. **(b)** Distribution of different types of *TERT* alterations across cancer types, as shown in Xena chart view (chi-squared, one-sided, F= 426.2.0, p<0.001). Xena automatically runs the appropriate statistical test for every chart.

UCSC Xena is best for users who are interested in integrating diverse types of PCAWG data from the same samples, such as simple mutation data, with gene expression and fusion calls, as well as more uncommon data types such as alternative splicing events, promoter usage, and mutational signature scores. It also provides a mechanism for viewing protected non-coding SNVs either separately or in conjunction with other PCAWG data. An example use case using Xena is found in Figure 2, demonstrating how a researcher can explore alterations in the TERT gene. Both public (structural variant data) and private data (SNVs) on the TERT gene is shown. Data is integrated in the browser, keeping private data protected. Even though the data is distributed across multiple hubs and those hubs have different access control, it appears to the user to come from a unified dataset, allowing easy visualization and data integration. In Figure 2 we see alterations by SNV and alterations by larger structural variation that are mutually exclusive. We also see that there is significant difference in the type of alteration in different cancer types (chi-squared, one-sided, F= 426.2.0, p<0.001).

### Chromothripsis Explorer and a Use Case

(http://compbio.med.harvard.edu/chromothripsis/). Chromothripsis refers to the mutational process characterized by massive *de novo* rearrangements that affect one or multiple chromosomes (Stephens 2011). The whole-genome data set assembled by PCAWG now provides an unparalleled opportunity to characterize chromothripsis patterns on a large-scale at single-base resolution across over 30 cancer types. Although statistical metrics are generally used to identify chromothripsis patterns (Korbel 2013), visual inspection still remains essential to dismiss false positive cases generated by other mechanisms of genome instability (Cortes-Ciriano 2018; Notta 2016). The Chromothripsis Explorer is an open source R Shiny application that visualizes chromothripsis patterns detected using WGS data (Cortes-Ciriano 2018; Campbell 2020).

The Chromothripsis Explorer provides tools for the exploration of chromothripsis frequencies and patterns across tumor types (Figure 3a). Specifically, it provides interactive Circos plots (Yu 2018) for each tumor, allowing researchers to explore large-scale alterations such as chromosome arm deletions, and complex mutational patterns such as chromothripsis and chromoplexy (Figure 3b). Each Circos plot is divided into 7 tracks that display, from outer to inner rings: (i) hg19 cytobands; (ii) inter-mutation distance and location for pathogenic (*i.e*., non-synonymous, stop-gain, and stop-loss) and nonpathogenic SNVs, and frame-shift and in-frame indels; (iii) chromothripsis regions; (iv) total copy number; (v) minor copy number profiles, defined as the least amplified allele, to visualize LOH regions; and (vi) gene track, and (vii) structural variations displayed according to the read orientations at the breakpoints (duplication-like SVs in blue, deletion-like SVs in orange, head-to-head in black, and tail-to-tail inversions in green). By hovering over the Circos plots, one can obtain information about a mutation of interest at single-base resolution, as well as gene annotations and functional effect predictions. In addition to genomics data, clinical and histo-pathological data are provided for all tumors in the form of customizable tables that permit to map tumor identifiers across cancer projects (*e.g*., TCGA to ICGC IDs; Figure 3b).

**Figure 3.**
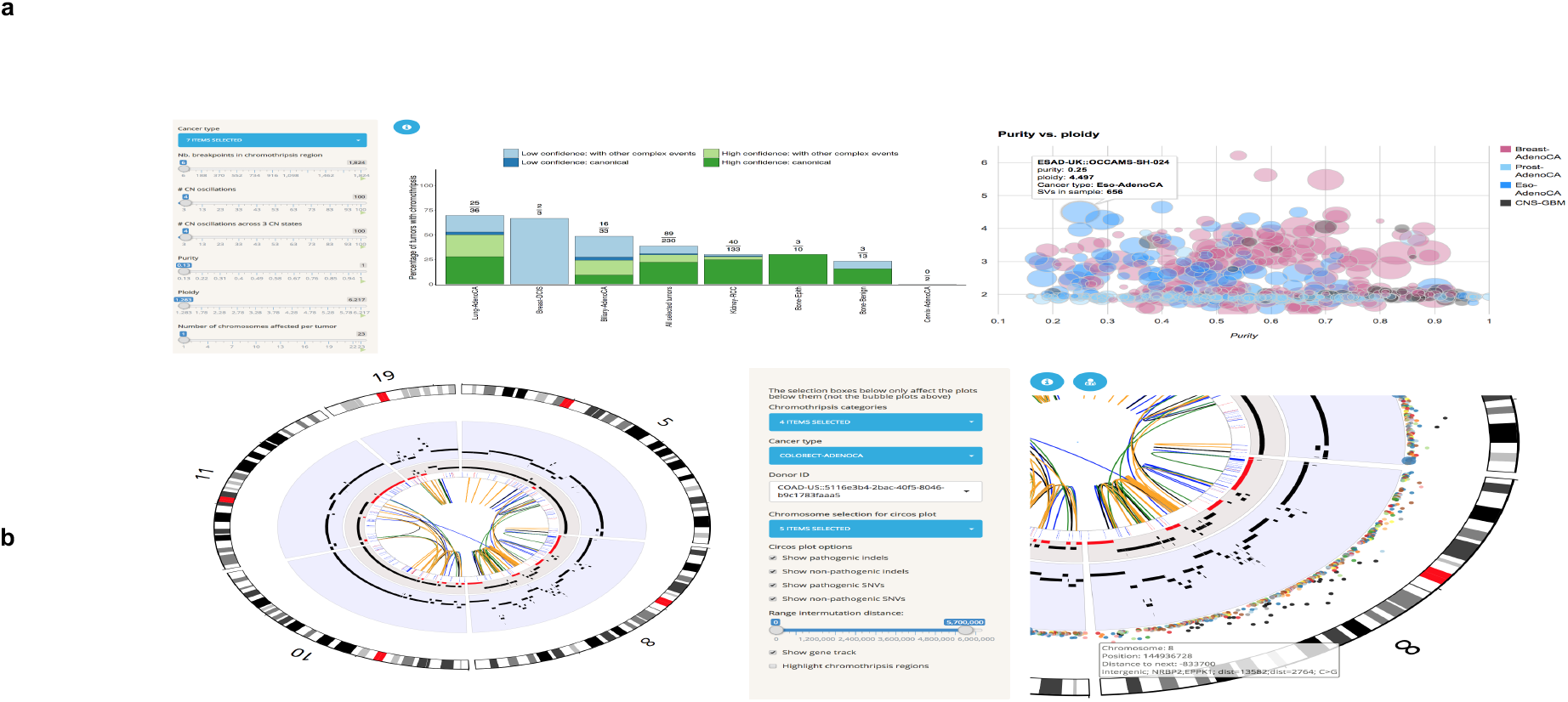
Functionalities of the Chromothripsis Explorer. **(a)** Interactive bar plot for the visualization of chromothripsis rates for selected cancer types. The left-hand side panel corresponds to variables used for the detection of chromothripsis patterns (e.g., number of copy number oscillations; Cortes-Ciriano 2018). The user can modify the values of these to explore chromothripsis rates across variable stringency criteria. The right-hand side panel illustrates additional functionalities to explore the relationship between purity and ploidy for tumors from selected cancer types. **(b)** Visualization of complex rearrangements involving 5 chromosomes in a ColoRect-AdenoCA patient (ICGC ID: DO9034). The right-hand panel shows a zoomed view of chromosome 8 that illustrates the tracks available in the Circos plots. From the outer to the inner ring, the tracks correspond to hg19 cytobands, SNVs (colored according to the mutation type and distributed according to the inter-mutation distance), total copy number over a blue background, minor copy number (LOH regions, i.e., with a minor copy number of equal to 0, are depicted in red), gene track, and SVs. Further information about the tracks can be accessed by clicking on the blue information circle located above the Circos plot.

The Chromothripsis Explorer is best for users who are looking for a global picture of the somatic alterations in a tumor, such as large-scale aneuploidies or translocations. It also provides visualizations of the point mutations, as well as small insertions and deletions, on a genome-wide scale. A representative use case for Chromothripsis Explorer is the exploration of complex rearrangements in human cancers, as shown for the ColoRect-AdenoCA tumor (ICGC ID: DO9034) depicted in Figure 3b. By selecting the chromosomes harboring massive rearrangements in this case chromosomes 5, 8, 10, 11 and 19, users can investigate the consequences of complex genomic rearrangements, such as the loss of heterozygosity across chromosome 8, and copy number amplifications.

### Expression Atlas and a Use Case

(https://www.ebi.ac.uk/gxa/experiments/E-MTAB-5200/). Expression Atlas is an added-value database and web-service that enables users to identify gene expression in different tissues, cell types, diseases and developmental stages. It collects, annotates, re-analyses and displays gene, transcript and protein expression data. It supports two types of study design: baseline and differential. Baseline studies involve quantifications of genes within tissues, developmental stages, cell lines, as well as others. Differential studies perform differential expression comparisons between different samples, for example disease versus healthy tissues (Figure 4). In addition to the PCAWG datasets, expression studies from archives such as ArrayExpress, GEO (Gene Expression Omnibus) and ENA (European Nucleotide Archive) are selected for further curation and processing. Data curation involves a semi-automated process of identifying the experimental factors, such as diseases or perturbations, annotating metadata with Experimental Factor Ontology terms (EFO) and describing the experimental comparisons for further processing. Currently, we provide results on over 3,500 experiments that include about 120,000 assays from over 60 different organisms. The data sets cover over 100 cell types from the Cell Ontology and over 700 diseases represented in the EFO.

**Figure 4.**
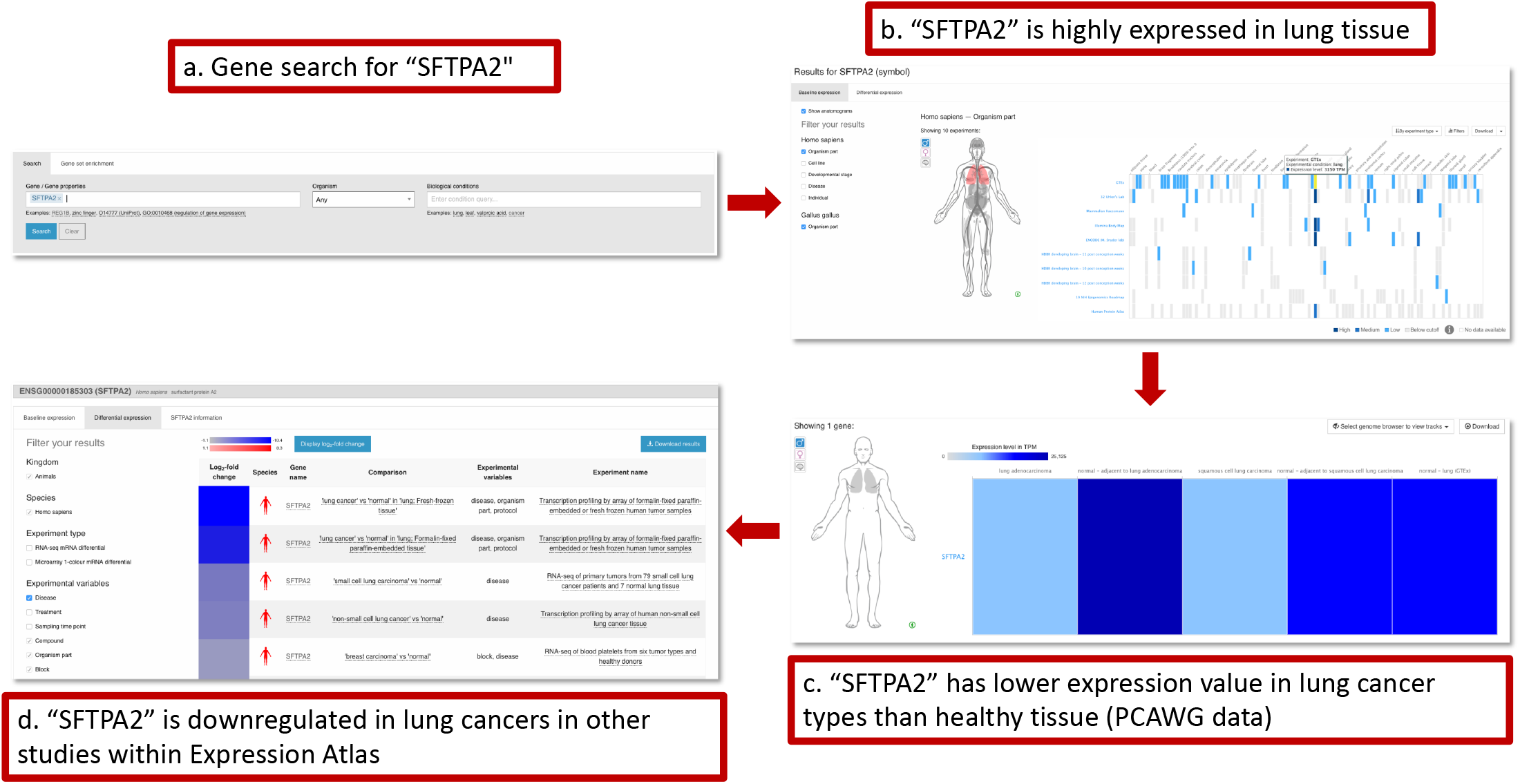
Example gene search in Expression Atlas. **(a)** Searching for experiments where *SFTPA2* is expressed or differentially expressed. **(b)** Viewing expression of *SFTPA2* in different tissues, across all baseline experiments, shows a consistent, high expression in lung. **(c)** Looking for the same gene in the PCAWG study within Expression Atlas, we see that it displays low expression in lung cancers (lung adenocarcinoma and squamous cell lung carcinoma), whereas it is highly expressed in normal tissue whether it is adjacent to lung adenocarcinoma or adjacent to squamous cell lung carcinoma. It is also highly expressed in lung samples from GTEx. **(d)** Finally, through the pages of differential assays in Expression Atlas, we confirm that *SFTPA2* is downregulated in further studies of lung cancer in Expression Atlas.

Expression Atlas includes large landmark baseline studies on human subjects or cell lines, such as GTEx, CCLE, ENCODE, BLUEPRINT, HipSci, as well as differential studies on human diseases in human patients or animal models. Analyses of bulk or single cell RNA-seq data sets are performed using our open source pipeline iRAP (Fonseca 2014). Expression Atlas can be searched by gene, gene set and experimental condition queries (Figure 4a). Gene, transcript and protein expression across different conditions is displayed through heatmaps and boxplots (Figure 4b). Annotation of data sets with EFO terms enables nested searching across related tissues, diseases and other conditions modeled within EFO. For example, when searching for “cancer”, the search will produce results for all different types of cancer, including “leukemia”. PCAWG datasets can be viewed and queried within their “study pages” or they can be viewed alongside other studies within Expression Atlas, returned as matches to gene or condition queries from the home page.

Expression Atlas is best for users who are interested in viewing how gene expression data from PCAWG compares with other resources, especially to ‘normal’ tissue in GTEx. It also provides the ability to see gene expression on an anatomical figure, making it easy to see patterns of expression across the body. An example use case of Expression Atlas in Figure 4 shows a typical gene search for gene *SFTPA2* to identify in which tissues it is expressed, and under which conditions its expression changes. The results of the query show high expression in lung tissue across different baseline expression studies available through Expression Atlas. Looking specifically into the PCAWG data sets, we see that it displays low expression in lung cancers (lung adenocarcinoma and squamous cell lung carcinoma), whereas it is highly expressed in normal tissue whether it is adjacent to lung adenocarcinoma or adjacent to squamous cell lung carcinoma. It is also highly expressed in lung samples from GTEx. Finally, through the panel of available differential studies (bulk RNA-Seq or microarrays) we can confirm that *SFTPA2* is down-regulated in further studies of lung cancer in Expression Atlas.

### PCAWG-Scout and a Use Case

(http://pcawgscout.bsc.es/). As opposed to only offering a limited and predefined list of analyses, PCAWG-Scout offers a variety of on-demand analysis functionalities. These analyses allow researchers to explore and visualize the data, form a hypothesis, run the relevant analysis, and immediately explore and visualize the results, giving rise to an analysis loop that drives discovery. These analyses are performed on data from the PCAWG main data release, as available in the ICGC data repository, and results from the different working groups. Results from PCAWG working groups include driver calls for different cohorts and for individual samples, mutation clonality assignments, and mutational signatures, which are all integrated into different sections of the PCAWG-Scout reports, tables, and interactive visualization graphics. The structure of the web interface accommodates reports for single entities (e.g. cohorts, samples, and genes) and lists of entities. These reports show relevant information about the entities, such as descriptions, statistics, plots, or interactive 3D protein representations and network graphs (Figure 5). Reports also offer further analyses that can be performed over the entities, such as enrichment analysis of gene lists, driver predictions over cohorts, survival analysis for lists of donors, or (non-authoritative) recommended therapies for individual donors (Supplementary Figure 4). The user can navigate these reports and analysis results as well as instantiate new reports as desired. The web interface uses a plugin approach to extend the reports and analysis that can be performed, making it easy for advanced users to add additional analyses. To help integrate PCAWG-Scout in a larger context, data and results are exported in interoperable formats.

**Figure 5.**
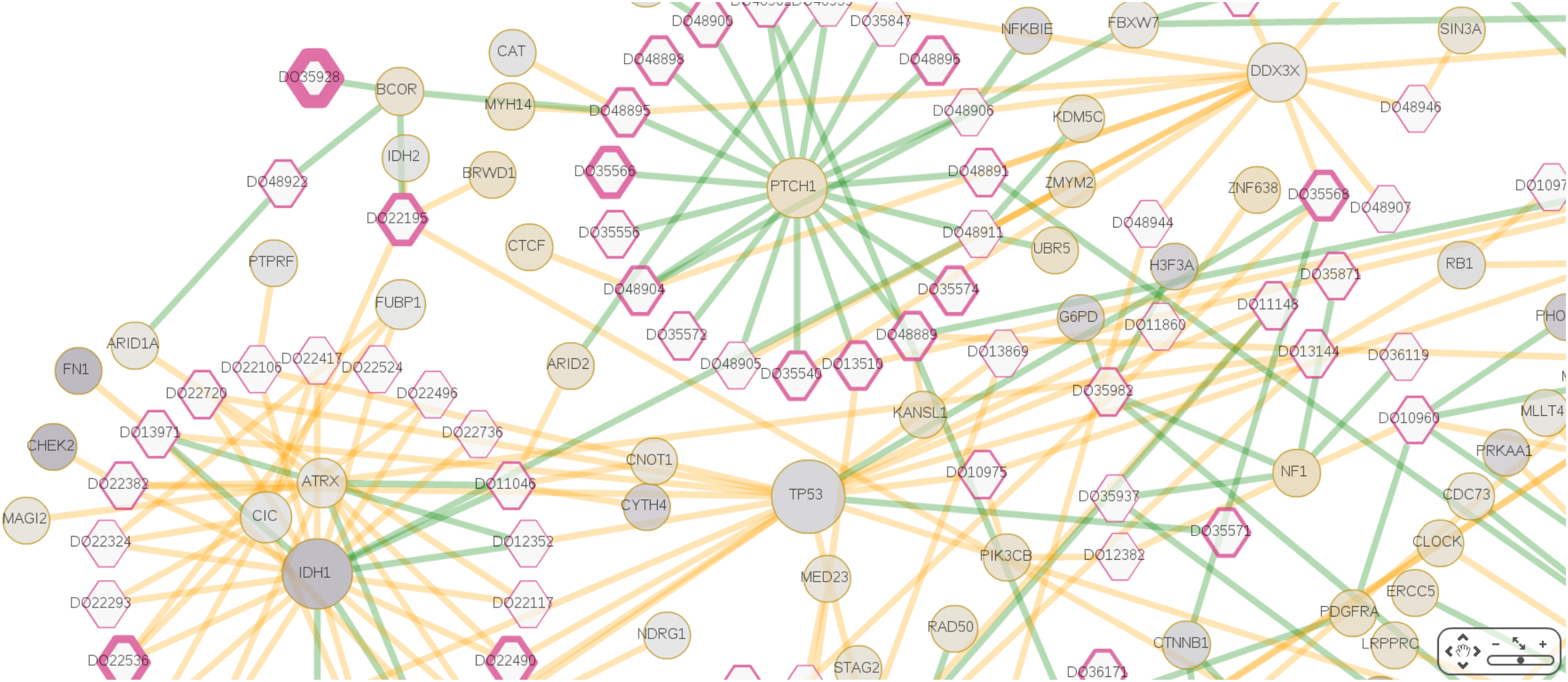
Customized visualization of the CNS meta-cohort donor driver events in PCAWG-Scout. This cytoscape-based visualization, available from the Study entity report page for the cohort, shows donors as hexagons and genes as circles. Edges represent driver events predicted (orange) and validated (green); donor driver events are curated by the drivers working group. Driver events for *IDH1* are mostly predicted while for *PTCH1* all are validated. The border size of the donors is their reported survival time. The size of the gene nodes is proportional to the significance of the extent to which mutations over these genes in this cohort are more damaging than expected (one-sided t-test comparing damage scores of mutations in cohort with all possible derived from SNV mutations, sample size varies with each protein); *IDH1*, *TP53* and *DDX3X* standout. The color of the genes is proportional to their differential expression t-value statistic when comparing tumor samples with *IDH1* mutations with the remaining tumor samples. The color is a gradient from repressed (dark purple) and overexpressed (gold) genes (using a two-sided t-test, FDR corrected, n=46), with an intermediate color for non-deregulated (light grey); *IDH1* is among the most repressed, along with *FN1* and *CHEK2*, which have driver events co-occurring with *IDH1* on some samples, while *PTCH1* is overexpressed on *IDH1* mutants. The extreme values for the gradient are taken genome wide and not limited to the genes present in the graph. Graphical aesthetics of node border-width, node size, node color, and edge color are configured interactively from data pulled from tables while exploring the network of linked reports around this cohort. Methods to reproduce this visualization are available in Supplementary Note 2.

PCAWG-Scout is best for users who are looking for a web interface to explore PCAWG data by running their own analyses, including differential gene expression. It also offers 3D mutation views for coding SNVs and INDELs. The potential to explore PCAWG data in PCAWG-Scout is illustrated in Figure 5. Here the network visualization tool was configured from the web interface using data gathered by running specific tools within. This plot offers the user a bird’s eye view of a number of important facets of the biology of central nervous system tumors. For instance, *IDH1*, *TP53*, and *DDX3X* also stand out as genes in which mutations are more damaging than expected. Plots such as these can help the user identify patterns which can then be further assessed on the web interface, such as mutual exclusivity, alternate means of deregulation of gene function by mutation or gene expression, or association with sample prognosis.

### Combining the strengths of each tool

While we can see that each tool has its strengths, using these tools together can help give a deeper understanding of tumor biology. We demonstrate this using a common driver event in prostate cancer: the fusion of the oncogene *ERG* (St. John 2012, Adamo 2016, Figure 1). Xena’s visual spreadsheet allows us to look across all 18 PCAWG prostate donor samples with both whole-genome sequencing and RNA-seq data and see that 8 of them harbor an *ERG* fusion. These donors also show *ERG* over-expression (Figure 1b). A view of the PCAWG structural variant data shows that across all donors, the fusion breakpoints are located at the start of *ERG*, leaving the *ERG* coding region intact while fusing to the promoter region of *TMPRSS2* or *SLC45A3* (Figure 1b). Additionally, we see that fusions detected by RNA-seq and whole-genome sequencing are not always consistent. Here, even using a consensus of detection methods (Campbell 2020), one fusion detected by a consensus of RNA-based detectors is missed, and the converse is also seen. This example shows that an integrated visualization across multiple data types and algorithms provides a more accurate model of a genomic event.

The Chromothripsis Explorer adds to this by showing a more in-depth view of the CNV and SV alterations in the 8 tumors with *ERG* fusions. In fact, we can see that the specific alterations for those 8 tumors vary widely (Figure 1c, Supplementary Figure 5). While donors DO36372, DO36359, DO36265 and DO36335 have quiescent genomes with few SVs, DO36356 and DO36283 show more complex karyotypes. For example, in tumor DO38283, chromosome 21 harbors multiple SVs that link it with chromosomes 2, 9, 13 and 21 (right). A closer look at the intrachromosomal SVs in chromosome 21 (left) reveals that the oncogenic fusion was generated by a deletion at chr21:39,988,805-40,578,907.

The Expression Atlas adds to this by showing how *TMPRSS2* and *SLC45A3* expression varies between PCAWG tumor samples and GTEx normal samples. In fact, we can see that both *TMPRSS2* and *SLC45A3* are highly expressed in normal prostate tissues and prostate tumors, as shown in the Expression Atlas Baseline Expression Widget (Figure 1d). Combined analysis of the PCAWG and GTEx datasets leads to the hypothesis that a subset of prostate cancers, through genome rearrangement, hijack the promoters of androgen-responsive genes to increase *ERG* expression, resulting in an androgen-dependent over-expression of *ERG*.

PCAWG-Scout adds to this by investigating genomic events in the samples that do not show ERG fusions. We know that although the *ERG* fusions are frequent, 46% (89 out of 195) of the PCAWG prostate tumors do not show them (Supplementary Figure 6). In fact, we can see using PCAWG-Scout’s mutual exclusivity analysis, that simple mutations in *FOXA1, SPOP, SYNE1*, and *ANKFN1* are significantly associated with these non-fusion tumors (Figure 1e). Furthermore, in PCAWG-Scout’s 3D protein structure view, the mutations in *SPOP* are shown to cluster tightly around the interaction interface for *PTEN* (Figure 1e), suggesting that those mutations may lead to altered protein function for *SPOP*.

This use case demonstrates each tool’s unique strengths and views and that together the five tools provide a clearer understanding of this genomic event, in this case into *ERG* fusions in prostate cancer. In this example we started with UCSC Xena, but it is possible to start with any of these tools and then investigate further.

### Discussion

The data generated by the PCAWG consortium is a resource for understanding complex cancer biology. Here we have described five tools that aim to put this resource into the hands of all researchers and incorporate outside genomics resources to help users gain insight into tumor biology. These tools are the ICGC Data Portal, UCSC Xena, Chromothripsis Explorer, Expression Atlas, and PCAWG-Scout, all of which are available from The PCAWG Data Portals and Visualization Page (http://docs.icgc.org/pcawg). While PCAWG data is rich in cancer biology insights, visualizing this data was challenging due to the large number of whole genomes. First, many visualization tools that focus on genes do not have a way to explicitly view the intergenic and intronic regions that whole genome sequencing elucidates. Second, the large size of whole-genome data imposes high performance requirements for interactive tools, especially those on the web. Finally, the analysis of whole-genome data provides an array of high-quality genomic information, including point mutations, gene fusions, promoter usage, and structural variants across thousands of samples and dozens of disease entities. Many visualization tools, especially those for users without extensive computational training, are currently limited to coding regions and more typical genomic datasets such as SNV or CNV. They are not equipped to be able to take advantage of depth and complexity of information at each point of the genome made available by the PCAWG consortium. Each of these tools presented here were either created or extended to address these whole-genome visualization challenges as part of the PCAWG project.

Despite these visualization challenges, another group from the PCAWG consortium has created an online tool to explore the panorama of driver mutations in PCAWG tumors and can be found via Gitools interactive heatmaps (http://www.gitools.org/pcawg), and browsed in IntOGen, at http://www.intogen.org/pcawg (Sabarinathan 2017). We envision that other tools will address the visualization challenges associated with whole-genome data that the PCAWG data will inspire new tools as well as be incorporated into other existing applications. We hope that this will be facilitated in part by the fact that the code for all our tools is open source and publicly available (Supplementary Table 3). We also made the functionality for some of the tools embeddable Javascript modules, such as for ICGC’s OncoGrid, UCSC Xena’s Visual Spreadsheet, and Expression Atlas’s Heatmap and Anatomogram Widget (Supplementary Table 4), so they can be incorporated into other 3rd party web applications.

## Acknowledgements

The ICGC Data Portal development is supported by the Ontario Institute for Cancer Research (OICR) through funding provided by the government of Ontario.

UCSC Xena development is supported by the National Cancer Institute of the National Institutes of Health under award numbers 5U24CA180951-04 and 5U24CA210974-02. The content is solely the responsibility of the authors and does not necessarily represent the official views of the National Institutes of Health.

Chromothripsis Explorer development is supported by the European Union’s Framework Programme For Research and Innovation Horizon 2020 (2014-2020) under the Marie Curie Sklodowska-Curie Grant Agreement No. 703543 (I.C.C.).

Expression Atlas development is supported by the European Molecular Biology Laboratory (EMBL) member states; the Single Cell Gene Expression Atlas from the Wellcome Trust (grant numbers 108437/Z/15/Z); the National Science Foundation of USA grant to Gramene database [NSF IOS #1127112]; Open Targets; Chan Zuckerberg Initiative.

PCAWG-Scout development is supported by joint BSC-IRB-CRG Program in Computational Biology and Severo Ochoa Award SEV 2015-0493.

